# phyloFlash – Rapid SSU rRNA profiling and targeted assembly from metagenomes

**DOI:** 10.1101/521922

**Authors:** Harald R. Gruber-Vodicka, Brandon K. B. Seah, Elmar Pruesse

**Affiliations:** Max Planck Institute for Marine Microbiology, Celsiusstraße 1, Bremen 28359, Germany; Department of Medicine, Division of Biomedical Informatics and Personalized Medicine, University of Colorado Denver Anschutz Medical Campus, Aurora, CO 80045, USA

## Abstract

The SSU rRNA gene is the key marker in molecular ecology for all domains of life, but is largely absent from metagenome-assembled genomes that often are the only resource available for environmental microbes. Here we present phyloFlash, a pipeline to overcome this gap with rapid, SSU rRNA-centered taxonomic classification, targeted assembly, and graph-based binning of full metagenomic assemblies. We show that a cleanup of artifacts is pivotal even with a curated reference database. With such a filtered database, the general-purpose mapper BBmap extracts SSU rRNA reads five times faster than the rRNA-specialized tool SortMeRNA with similar sensitivity and higher selectivity on simulated metagenomes. Reference-based targeted assemblers yielded either highly fragmented assemblies or high levels of chimerism, so we employ the general-purpose genomic assembler SPAdes. Our optimized implementation is independent of reference database composition and has satisfactory levels of chimera formation. Using the phyloFlash workflow we could recover the first complete genomes of several enigmatic taxa, including Marinamargulisbacteria from surface ocean seawater. phyloFlash quickly processes Illumina (meta)genomic data, is straightforward to use, even as part of high-throughput quality control, and has user-friendly output reports. The software is available at https://github.com/HRGV/phyloFlash (GPL3 license) and is documented with an online manual.

## INTRODUCTION

Shotgun metagenomics is a powerful tool to explore the functions of microbial communities, and to determine their phylogenetic or taxonomic composition (1, 2). It can avoid biases and artifacts associated with PCR-based amplicon methods, such as primer bias (3) and sequence chimerism (4, 5). Shotgun sequencing also yields more information because it samples from whole genomes rather than a single marker. Various approaches have been applied to profile the composition of shotgun metagenomic libraries. A common choice is local alignment against reference sequences, which may target a specific region in a single gene, (6, 7), curated sets of conserved single-copy genes (8, 9), or whole-genome data (10, 11). Others use alignment-free methods such as k-mer matching to classify reads (12–14), and some tools also go beyond classification to perform targeted assembly of specific genes (15, 16). Regardless of the method that is used for taxonomic or functional profiling, the results will be limited by the reference data available. For example, target organisms or their close relatives may not yet be represented in the database, and horizontal gene transfer can result in conflicting phylogenetic signal, especially in prokaryotic microbes (17, 18).

In molecular ecology, the gene for the small subunit ribosomal RNA (SSU rRNA) is the most important marker because it can be readily used to link sequences to actual cells. The SSU rRNA is available in high copy numbers in ribosomes and can be accessed through the well-established molecular probing technique of fluorescence *in situ* hybridization (FISH) (19, 20) Its value in imaging based analyses together with its high phylogenetic reliability due to low rates of horizontal transfer has made the SSU rRNA gene the best-sampled marker in terms of phylogenetic diversity (21). Even with the current advances of metagenomics, where drafts of microbial genomes can automatically extracted (“binned”) from metagenome assemblies (22), the SSU rRNA gene has the same essential role and is crucial for phylogenetics, imaging, and experimental verification. However, despite all the progress on automated binning (23), most metagenome assembled genomes (MAGs) do not even contain fragments of the SSU rRNA gene, not to mention the full gene (1, 24).

Ideally, we would like to leverage the vast existing knowledge base of the SSU rRNA gene in (meta-)genomics projects for several different outcomes: taxonomic profiling without assembly (6, 7), targeted assembly of full-length sequences for phylogenetics and probe design (15, 16, 25, 26), and linking SSU rRNA sequences to complete genomes (27). For each of these aims, separate software tools have already been developed, each with their own merits and shortcomings (16). However, these are ultimately different aspects of the same problem and should be considered together.

Here we describe phyloFlash, an open-source pipeline for the rapid profiling and targeted assembly of SSU rRNA from metagenomes. We use the pipeline to evaluate the trade-offs in different steps of taxonomic profiling and targeted assembly, starting from the preparation of the reference database, to the extraction and assembly of target reads, and finally to linking the assembled sequences to genome bins. We compare the performance of rRNA-specialized vs. general-purpose read mappers and assemblers, by using both simulated read data and real-world environmental metagenomes. We also reanalyze published data to show how the pipeline can be used to assess and compare the composition of metagenomes. Finally, we show how genome bins from metagenomes can be linked to SSU rRNA sequences, to bridge the divide between genome-centric metagenomics and experimental molecular ecology.

## MATERIAL AND METHODS

### Implementation of phyloFlash

The phyloFlash pipeline is written in Perl 5, with additional scripts in Perl and R for downstream analyses such as visualizations or multi-sample comparisons. The pipeline is outlined in Figure 1, and a manual is available at https://hrgv.github.io/phyloFlash/.

**Figure 1.**
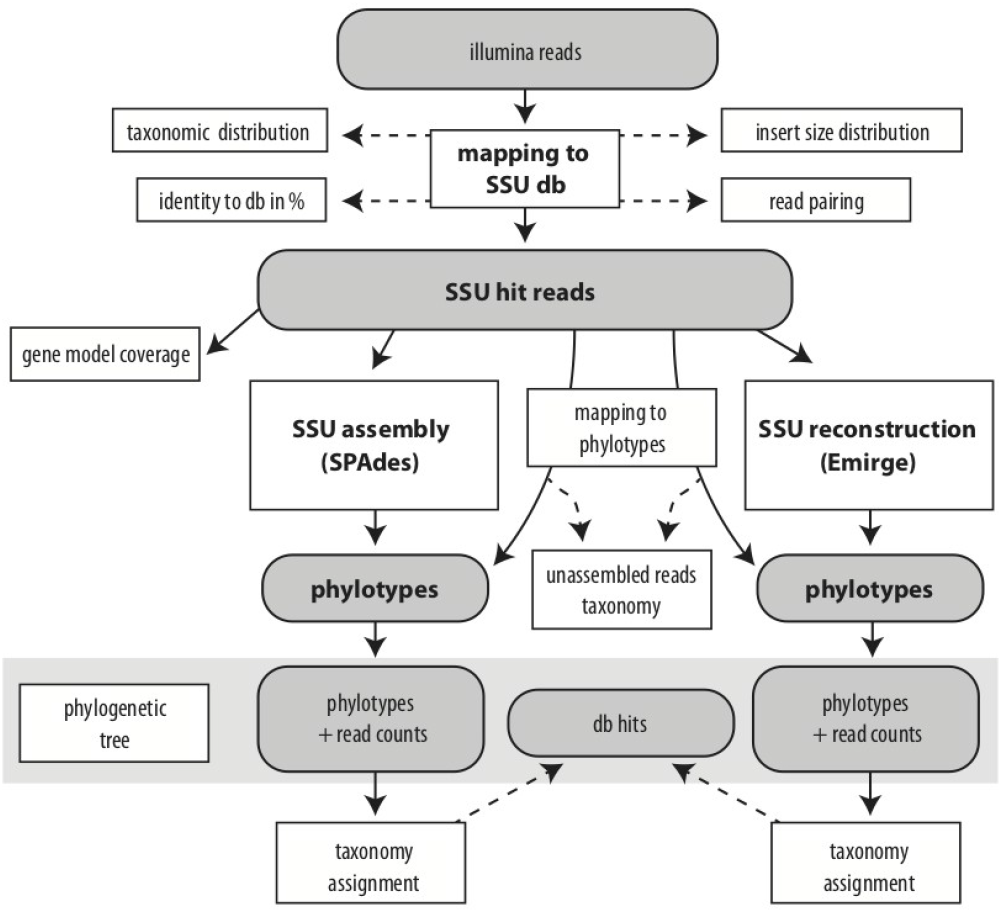
Flowchart of the phyloFlash pipeline. Grey boxes depict main data files, white boxes with bold print show processes. Full arrows indicate primary workflow, dashed lines processing of secondary output.

#### Database filtering and preparation

The pipeline uses the SSU Ref NR99 database from the SILVA project (https://www.arb-silva.de) (28). Sequences containing fragments of the large subunit (LSU) rRNA are detected with a hidden Markov model (HMM) from a customized version of Barrnap (E-value cutoff 10^−10^, >10% of total model length) (https://github.com/tseemann/barrnap), and are removed. Sequence regions containing low-complexity sequence or repeat k-mers are masked with bbmask.sh (from BBmap, https://sourceforge.net/projects/bbmap/) using k-mer lengths between 4 and 8 to detect both repeats and low-complexity, with a minimum masked sequence length of 20 bp, and entropy cutoff of 0.7. Known sequencing or cloning vectors are filtered by matching against the NCBI Univec database (ftp://ftp.ncbi.nlm.nih.gov/pub/UniVec/) with bbduk.sh (BBmap), using k-mer length 27 (down to 11 at ends of sequences), maximum Hamming distance of 1, and discarding sequences below 800 bp after trimming. The sequences are converted from RNA to DNA alphabet, and ambiguous bases are replaced by random base characters. The sequences are then clustered with VSEARCH cluster_fast at 99% (“NR99”) and 96% identity (“NR96”) (29). The NR99 sequences are indexed for BBmap with bbmap.sh, and in UDB format with VSEARCH (v2.5.0+). The NR96 sequences are indexed for Emirge (15) with Bowtie (30), and for SortMeRNA (31) with indexdb_rna (from SortMeRNA). The script phyloFlash_makedb.pl automates the database preparation steps described above. phyloFlash_makedb.pl can be used to update to new releases of the SILVA database and has been tested with the latest available SILVA release (132 at the time of submission) *SSU rRNA read extraction*. The inputs for phyloFlash are shotgun metagenomic paired-end libraries, which have been generated by an Illumina sequencer. SSU rRNA reads can be extracted either with BBmap or SortMeRNA.

For BBmap, the input reads are aligned (mapped) against the filtered NR99 database, with minimum identity 70% by default, retaining all ambiguous alignments if there are multiple best-scoring hits. Insert-size and mapping-identity histograms are also reported to evaluate library quality. Output is written in SAM and Fastq formats, retaining all read pairs where at least one read could be aligned. Known limitations in the BBmap implementation of SAM bitwise flags and read name handling are addressed.

For SortMeRNA, the input reads are first reformatted to uncompressed interleaved Fastq format with reformat.sh (BBmap). The reads are then aligned against the filtered NR96 database, using min_lis of 10 and E-value cutoff 10^−5^, reporting the 10 best alignments. Aligned reads are reported in SAM and Fastq formats. The SAM file is further processed to edit the bitwise flags, as SortMeRNA does not report complete pairing information in its implementation of the SAM format.

#### Taxonomic summary from read mapping

The edited SAM files from the read extraction step are read into memory. For each read pair, the taxonomic affiliation is assigned by taking the last common ancestor (LCA) of the taxonomy strings of all the database hits, using the SILVA taxonomy. The counts of LCA-consensus taxa for the library are then summarized at a user-specified taxonomic level between Domain and Species (Class by default).

#### Assembly of full-length SSU rRNA sequences

The extracted reads are assembled to full-length sequences using SPAdes (32) and optionally with Emirge.

For SPAdes, k-mer lengths are chosen based on the input read lengths. For reads ≥ 134 bp, k-mers of length 99, 111, and 127 bp are used. For shorter reads, the k-mer lengths used are next lower odd number to the read length minus 27, 17, and 7 bp. After assembly, contigs are screened for SSU rRNA sequences with HMM models for Bacteria, Archaea, and Eukaryota as implemented in the customized version of Barrnap. SSU rRNA regions under the E-value cutoff of 10^−100^ and at least 0.6 times the full-length model are extracted with Bedtools (33).

For Emirge, read pairs are run as single reads when their length is > 152 bp. For paired-end input, the average insert size estimated from the initial mapping with BBmap is used if the insert size is greater than 2.2 times the read length, otherwise a minimum insert size of 2.2 times the read length plus 0.5 is used. Emirge is run with 40 iterations.

To estimate the proportion of the extracted reads that were successfully assembled, the reads are re-mapped to the assembled sequences with bbmap.sh (BBmap) in fast mode using minimum identity of 98%. The closest-matching database sequences are also found with usearch_global (34) as implemented in VSEARCH at minimum identity 70%. Assembled sequences and their closest database hits are then aligned with Mafft (35) to produce an alignment and guide tree.

#### Reporting of results

Results are reported in both plain-text and HTML formats; the HTML report features interactive SVG-formatted graphics and allows the results to be shared easily in a single file that is compatible with modern web browsers without additional software. The report contains the taxonomic summary, mapping statistics, tree and tables of assembled full-length sequences and their closest database hits. All outputs, including mapping files and assembled sequences, can also be compressed into a single archive file at the end of the phyloFlash run.

#### Comparison of multiple samples

The phyloFlash results of multiple metagenomes can be graphically compared with the phyloFlash_compare.pl script. This uses the mapping-based taxonomic summary to generate either a barplot or heatmap of taxa for each library, at a user-specified taxonomic level. The clustering of libraries in the heatmap can either treat each taxon as independent, or account for the taxonomy in calculating the distance matrix, using a Unifrac-like metric.

The abundance-weighted taxonomic Unifrac-like metric treats the hierarchical taxonomy as a tree where each taxon is represented by a node *i* associated with a branch of unit length. The raw weighted Unifrac value *u* of (36) therefore reduces to 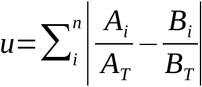 where *n* is the total number of nodes in the tree, *A_i_* and *B_i_* are the numbers of sequences in samples A and B respectively classified to node *i* or its descendants, and *A_T_* and *B_T_* are the total numbers of sequences in samples A and B respectively. Furthermore, as the taxonomy tree is ultrametric (all branches of the same length), the parameter *d_j_* of (36) reduces to the taxonomic level at which the comparison is performed, e.g. 4 for ‘Order’ in the Linnaean hierarchy. Then the scaling factor *D* = 2*d* where *d* is the taxonomic level, and the normalized abundance-weighted taxonomic Unifrac-like metric is 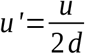.

### Evaluation of unfiltered vs. filtered database

#### Database preparation and indexing

The SILVA release 132 SSU Ref NR99 database was downloaded from arb-silva.de in Fasta format. For the unfiltered database, sequences were clustered at 96% identity with VSEARCH v2.5.0 (“NR96”) and converted from RNA to DNA alphabet, and any ambiguous bases were replaced by random base characters. Indices were built for BBmap v37.99 and SortMeRNA v2.1b with default parameters. The filtered database was prepared as described above for the phyloFlash pipeline. Read mapping for this comparison was performed with E-value cutoff 10^−5^ for SortMeRNA (this is the default cutoff used by the Matam assembler) and minimum mapping identity 70% for BBmap on 10 metagenomic libraries from the Tara Oceans project, using only the first 10 million read pairs per library (Supplementary Table 3) (37).

#### Calculation of sequence entropy and redundancy

The number of possible DNA k-mers of length *k* is *N* = 4^*k*^. For a sequence of length *L*, k-mers are counted by applying a sliding window of width *k* across the sequence. The total occurrences of a given k-mer *i* is *w_i_*. The total count is 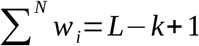. The probability of each k-mer is 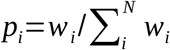. Hence the entropy of the sequence using k-mer length *k* is 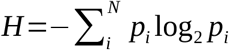.

The redundancy *R*=1 −*H*/log_2_ *A*, where *A* = *N* if *L*−*k* +1 >*N* and *A* = *L*−*k* + 1 otherwise.

This is because if *L* is short and *k* is long, the maximum possible entropy of the sequence is constrained by the length of the sequence. Redundancy is thus always between 0 (maximum entropy) and 1 (zero entropy). The calculation was performed on the SAM file from each mapping with the script readEntropy.pl (https://github.com/kbseah/misc_tools/).

### Evaluation of read-mapping and assembly methods

#### Simulated metagenome

Shotgun metagenome paired-end reads were simulated from 10 bacterial RefSeq genomes (Supplementary Table 1) with the software package ART v2.5.8 (38) at average 50-fold coverage per genome, with the first library using the Illumina HiSeq2000 error profile (100 bp read length, insert size 220 ± 110 bp), and the second library using a HiSeq2500 profile (150 bp, insert size 300 ± 110 bp). Plasmid sequences were excluded from the simulation. To compare the effect of different mapping settings on read extraction, the phyloFlash pipeline (v3.3) was run with either BBmap (at three different minimum identity cutoffs: 50%, 60%, and 70%) or with SortMeRNA (at three different E-value cutoffs: 10^−5^, 10^−7^, 10^−9^), and using the filtered SILVA 132 SSU Ref database as described above (“Implementation of phyloFlash pipeline”). Overlap between the reads aligned by each method was calculated with the script samDiff.pl (https://github.com/kbseah/misc_tools/). Coverage of genomic features was calculated with featureCounts v1.5.2 using the RefSeq feature tables and the SAM-formatted alignment produced by ART to track the extracted reads back to the original genomes. Timings for BBmap and SortMeRNA read extraction given in Supplementary Table 2 were based on runs on the same computer using eight processors (Intel Xeon CPU E5-1650 v2, 3.5 GHz).

SPAdes v3.11.1 and Emirge v0.60 were used to assemble extracted reads within the phyloFlash pipeline, using the parameters described above. The library was also separately assembled with Matam v1.4.0 (16) for comparison, using the filtered SILVA 132 SSURef NR96 database, extracting reads with SortMeRNA at E-value 10^−5^, and with other parameters at default. The scaffolds assembled by each tool from the read set extracted by SortMeRNA at E-value 10^−5^ were aligned against SSU rRNA sequences identified from the original genome assemblies by Barrnap, using Mafft v7.187 (FFT-NS-2 strategy) (35). The phylogeny was inferred from the alignment with FastTree v2.1.7 (39) using the Jukes-Cantor substitution model, and CAT approximation with 20 rate categories.

#### Environmental metagenomes

85 metagenomic libraries from the Tara Oceans project (37) were used to evaluate performance on real-world data (Supplementary Table 3). For each library, only the first 10 million read pairs were used, to control for the effect of library size on assembly quality. Different mapping settings were tested as described above. Assembly with SPAdes, Emirge, and Matam was performed as described above. The uchime_ref chimerism score (5) was calculated for each assembled SSU rRNA gene sequence with VSEARCH v2.5.0, using the filtered SILVA 132 SSURef NR99 database as reference.

### Usage examples

#### Low-diversity metagenome and reference database completeness

The metagenomic library (accession PRJEB29979) of a small marine flatworm and its associated symbiotic bacterium was processed with phyloFlash with the option -everything (BBmap mapper, SPAdes and Emirge assemblers), using either the default SILVA 132 SSURef database or version of the database where all sequences with ≥87% identity to the flatworm and symbiont SSU rRNA genes were removed. Sequences were also assembled with Matam as described above for the simulated metagenome. The library was also processed with Kaiju v1.4.5 (13) using the NCBI nr protein database prepared with the Kaiju script makeDB.sh −e (prokaryotes and microbial eukaryotes), in “greedy” mode allowing up to 5 mismatches. Kaiju was run again using only reads mapping to the complete symbiont bacterial genome (see below for the binning procedure), which were extracted with bbmap.sh (“fast” mode). The Kaiju output was summarized at the taxonomic class level.

#### Comparison of multiple environmental metagenomes

Publicly available metagenomic libraries from seven gutless oligochaete worm species, whose symbiotic bacteria had previously been characterized by Sanger-sequenced clone libraries of PCR-amplified SSU rRNA (Supplementary Table 5), were processed with phyloFlash with the option -everything. Samples were compared as a heatmap on the basis of mapping-based taxonomic composition at the order level using phyloFlash_compare.pl with options --task heatmap,barplot --level 4.

#### SSU rRNA-based binning of microbial genomes

Libraries from the simulated metagenome (150 bp), *Paracatenula* and gutless oligochaete animal metagenomes, and selected Tara Oceans metagenomes were assembled for the SSU rRNA-targeted genome binning example (Supplementary Table 6). Each assembly was performed with the same protocol, using both the Megahit v1.1.3 (40) and MetaSPAdes v3.12.0 (32) assemblers (code in Supplementary Information). Each read library was first screened with phyloFlash (option -everything). Sequencing adapters and low-quality bases (minimum Phred score 2) were trimmed from both ends of the reads with bbduk.sh (from BBmap suite). Reads with k-mer depth below 6 (k=31) were filtered out with bbnorm.sh (BBmap). The trimmed, high-k-mer-coverage reads were assembled with Megahit using minimum k-mer length 21, maximum k-mer 141, with steps of 20; the Fastg file was generated with megahit_toolkit contig2fastg from the k=141 contigs, and binned with phyloFlash_fastgFishing.pl, keeping only SSU rRNA above 60% full-length and contig clusters above 100 kbp total length. The reads were also assembled with MetaSPAdes with k-mer lengths 21,33,55,77,99,127, and binned in the same manner. Bin quality (completeness, contamination) was estimated with CheckM v1.0.11 using the taxonomy workflow and Bacteria domain marker set.

## RESULTS AND DISCUSSION

### Reference database artifacts influence targeted read extraction

The preparation of the reference database affects the number and quality of reads extracted for downstream analysis, even when using a curated resource such as the SILVA databases. Although potential contaminants and low-complexity sequences represented only a small fraction of the total sequences, they had a disproportionate effect on the reads recovered. In release 132 of the SILVA SSU Ref NR99 database (total 695171 sequences, 1.006 Gbp), we detected 42.88 kbp (in 151 sequences) of matches to partial LSU rRNA, and 1.102 Mbp (in 56075 sequences) of potential vector contamination. Furthermore, sequence-repeat and low-complexity regions constituted 23.31 kbp.

Fewer reads were extracted with a filtered database, but their quality was improved. For example, in library ERR315856, 32937 reads were recovered by SortMeRNA (E-value 10^−5^) before filtering, but only 17227 afterwards. Moreover, the mean informational redundancy decreased from 0.0748 to 0.0151, indicating that fewer low-complexity sequences were extracted (Figure 2). The degree of impact from filtering depends on the metagenome library composition, but similar results were observed when reads were extracted with BBmap, and for other metagenome libraries (Supplementary Figure 1). Subsequent results reported here used the filtered database.

**Figure 2.**
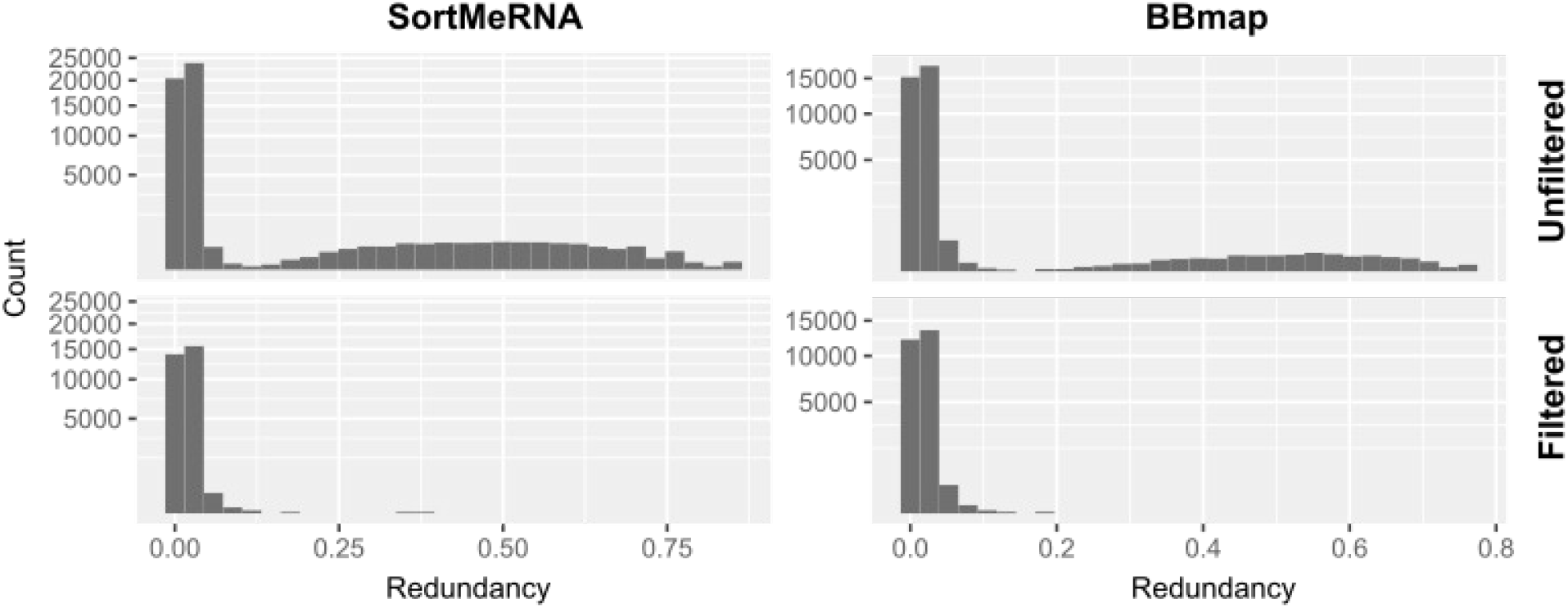
Database filtering removes low complexity read hits. Informational redundancy (k=5) in reads extracted by SortMeRNA (left) or BBmap (right) using unfiltered (above) and filtered (below) reference databases, for library ERR315856.

Some SSU rRNA genes may indeed contain low-complexity regions, nonetheless it is still desirable to remove them from the reference database. This is because any low-complexity sequence in the library (e.g. from eukaryotic genomes), regardless of origin, will tend to map to these regions. This would skew any analysis of the taxonomic composition of a metagenome that is based on mapping to reference sequences, and could also affect assembly and other downstream analyses.

### BBmap provides similar sensitivity and higher selectivity vs. SortMeRNA at five-fold faster runtimes

Different tools have been developed for extracting SSU rRNA reads from shotgun metagenomes. Those that depend on alignment to a reference database (“mapping”) can be used to derive an approximate taxonomic assignment from the closest-matching reference sequences, which is not possible with model-based methods (e.g. HMMs or covariance models). We compared two reference-based tools for read extraction: SortMeRNA, originally designed to identify rRNA reads in metatranscriptomes (31), and BBmap, a general-purpose mapper.

The performance of these read extraction tools was dependent on their stringency settings (E-value cutoff for SortMeRNA, minimum mapping identity for BBmap), as well as properties of the read library such as read length, insert size (for paired end reads), and error profile. We simulated two Illumina paired-end metagenomes from the same ten RefSeq bacterial genomes (Supplementary Table 1): a HiSeq2000 library with 100 bp reads (220 ± 110 bp insert size, mean ± standard deviation) and a HiSeq2500 library with 150 bp reads (300 ± 110 bp insert size). In terms of their *sensitivity*, i.e. the fraction of reads originating from annotated SSU rRNA features that they could recover, both tools had a similar performance (ca. 94–96%, depending on settings) with 100 bp reads, and SortMeRNA performed slightly better (96–97%) than BBmap (93–96%) with 150 bp reads (Figure 3, Supplementary Table 2). However, in terms of *selectivity*, i.e. the proportion of true SSU rRNA reads among the total reads recovered, BBmap performed better with 150 bp reads than 100 bp reads (86–95% vs. 82–92%) whereas SortMeRNA performed worse (73–82% vs. 82–89%). The main source of false positive hits were tRNA genes (Figure 3). BBmap was also about five-fold faster than SortMeRNA when run on the same machine (Supplementary Table 2).

**Figure 3.**
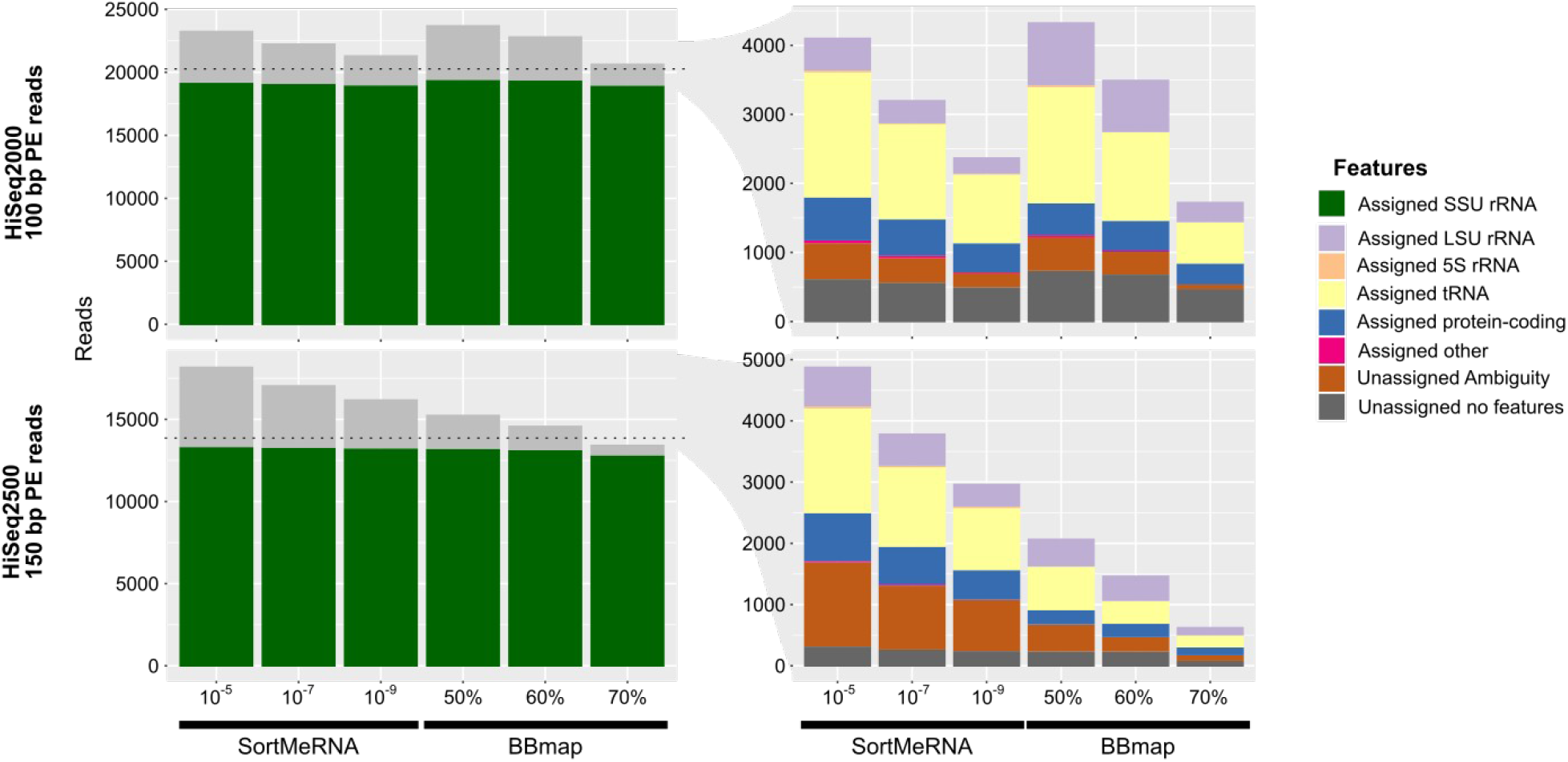
False positive read extraction depends on stringency cutoffs, but only BBmap results improve with read length. Origin of reads extracted by SortMeRNA and BBmap at different parameters (horizontal axes) from simulated metagenome of 100 bp (above) or 150 bp reads (below), showing the proportion of total originating from actual SSU rRNA features (left), and the origin of non-SSU rRNA extracted reads (right). Horizontal lines indicate the total actual SSU rRNA-origin reads in each library (20213 and 13829 respectively).

### SortMeRNA can yield more assembled reads than BBmap with environmental metagenomes

In real metagenomic libraries, the origin of each read is not known *a priori*. The extraction of SSU rRNA reads can only be evaluated indirectly, by comparing the overlap in reads recovered by both tools. An alternative measure for the quality of read extraction is the fraction of extracted reads that can be assembled to full-length gene sequences. We performed both comparisons with 85 environmental metagenomes from the surface ocean seawater habitat (Supplementary Table 3).

More restrictive settings (lower E-value cutoff for SortMeRNA, higher minimum identity for BBmap) resulted in more overlap between the reads recovered by both tools, with a mean 87.2% overlap for the most restrictive settings tested (SortMeRNA E-value 10^−9^, BBmap 70% min. identity) vs. a mean 66.6% overlap for the least restrictive settings tested (10^−5^, 50%) (Figure 4). More permissive settings for a given tool resulted in more reads extracted only by that tool (Supplementary Figure 2). Similarly, more restrictive settings resulted in a higher fraction of reads that could be assembled into full-length SSU rRNA sequences, e.g. for SortMeRNA, the mean fraction was 45.4% at E-value cutoff 10^−9^ vs. 40.6% at E-value 10^−5^, and for BBmap, mean 41.1% at minimum mapping identity 70% vs. 34.2% at >50% minimum identity. This can be seen in Figure 4b as the slope of the relationship between the number of assembled reads vs. the total extracted reads increased as settings became more restrictive, irrespective of the extraction tool.

**Figure 4.**
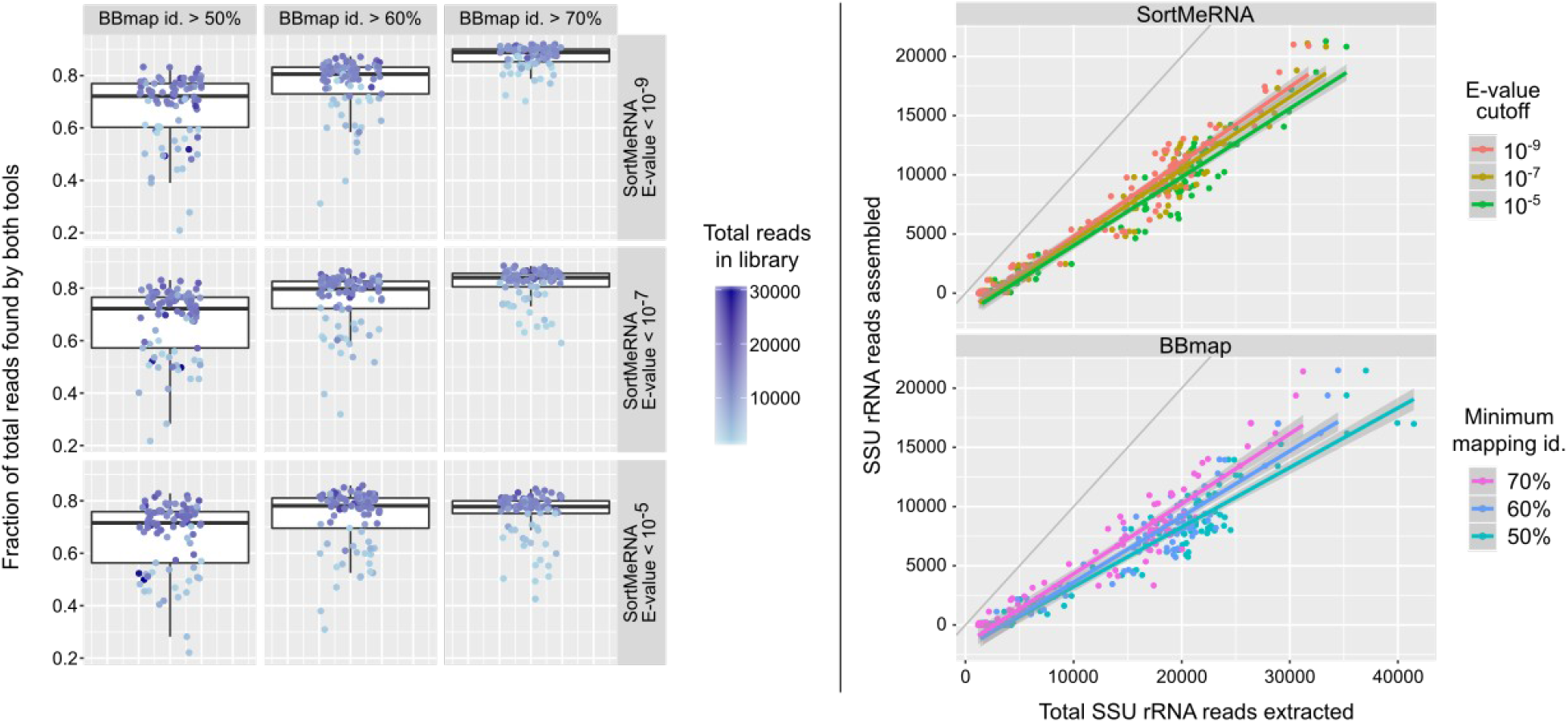
Mapper and cutoff settings influence SSU rRNA reads extraction from Tara Oceans metagenome libraries but yield highly similar assembly ratios. (a) Fraction of reads extracted by both tools at different BBmap minimum identities (horizontal) and SortMeRNA E-value cutoffs (vertical). (b) Reads assembled by SPAdes vs. total reads extracted for different parameter settings (colors) of SortMeRNA (above) and BBmap (below), overlaid with linear regression lines and 1:1 line for reference (in grey).

The absolute number of reads that can be assembled does not vary much for most libraries with the cutoff settings, and on average, SortMeRNA performs better than BBmap in extracting reads that can be assembled, with this improvement being more pronounced for some libraries (Supplementary Figure 3). The mean ratio of assembled reads extracted by BBmap to SortMeRNA is between 85.9% to 90.6% depending on the program settings being compared (Supplementary Table 4), but for more than half the libraries, both BBmap and SortMeRNA have comparable performance: out of the 81 libraries with a nonzero number of assembled reads, the BBmap:SortMeRNA ratio was >90% for 47 libraries, and >95% for 37 libraries.

We chose BBmap as the default read extraction tool as it was much faster in our tests with both simulated and real metagenomes and had similar performance in most cases. For more detailed but time-intensive analyses, and to investigate the possibility of highly-divergent taxa by targeted assembly, both SortMeRNA and more permissive settings for both tools can be tested. For BBmap, this comes with the caveat that the taxonomic summary would likely become less reliable. More permissive settings for initial read extraction may increase the probability of false positives, but can also make it possible to detect possible distant relatives. Given the popularity of SSU rRNA as a phylogenetic marker, and that SSU rRNA sequences in public databases represent the accumulated results of several decades of taxon sampling, the more restrictive settings should be adequate for the routine assessment of taxonomic composition using read-mapping hits to reference sequences. Defaults of E-value < 10^−9^ for SortMeRNA and minimum identity >70% for BBmap were therefore chosen for the phyloFlash pipeline.

### General-purpose assembler SPAdes yields longer and more divergent scaffolds at acceptable levels of chimerism

Existing software tools that have been developed for the targeted assembly of SSU rRNA sequences employ various algorithms and reference data (Table 1). The ideal assembler should have the following properties: be able to assemble full-length gene sequences, while avoiding chimeric assemblies, be able to assemble highly-divergent sequences, and yet differentiate between closely-related strains or paralogs. In reality, optimizing for one metric may preclude another. For example, attempting to recover strain-level variation will result in more fragmented assemblies, because insert sizes from Illumina shotgun libraries are usually too short to resolve full-length sequences that only differ in the variable regions of the SSU rRNA gene.

**Table 1.** Comparison of targeted assembler software for SSU rRNA sequences. * as of 2018-10-08

| Tool | Read extraction | Data-base | Method | Taxon classifier | Availability | Documentation | License | Last update* |
| --- | --- | --- | --- | --- | --- | --- | --- | --- |
| Emirge | Bowtie (v1) | SILVA | Expectation-maximization | (From mapping) | GitHub | Readme, User forum | GPL3 | 2016-11-09 |
| REAGO | Infernal (CM) | - | Overlap graph, pruning, paired-end guided path-finding | - | GitHub | Readme | BSD | 2017-01-12 |
| Matam | SortMeRNA | SILVA NR95 | Overlap graph, compression, SGA assembler on subgraphs | RDP classifier | GitHub, Conda, Docker | Readme | AGPL3 | 2018-09-11 |
| RAMBL | Bowtie2 | Greengenes | Taxonomic tree search and Dirichlet process clustering | RDP classifier | GitHub | Readme | Not specified | 2017-03-18 |

We compared three different assemblers, two targeted-assembly tools Emirge (15) and Matam (16) and the general-purpose genome assembler SPAdes (32), on SSU rRNA reads extracted with SortMeRNA using the filtered SILVA 132 SSURef NR96 reference database and E-value cutoff 10^−5^. To ensure a uniform comparison, we used only sequences matching at least 60% of a full-length SSU rRNA HMM model.

We first tested the assemblers on the same simulated metagenome previously used to test read extraction. This dataset originally contained a total of 25 SSU rRNA gene copies from ten species, which reduced to eleven sequences when clustered at 99% identity (Supplementary Table 1). SPAdes assembled the same ten scaffolds (one per genome) each time from both the 100 and 150 bp libraries, whereas the scaffolds assembled by Emirge and Matam differed between the two libraries (Supplementary Figure 4). Although Emirge also assembled ten scaffolds from each library, some were chimeric sequences that did not have a close match to any of the original sequences. Scaffolds assembled by Matam were often fragmented, e.g. from the 100 bp library, full-length sequences were not assembled for four genomes.

Some organisms have multiple SSU rRNA gene copies which may be only slightly divergent and pose a challenge to assembly. Included in the simulated metagenome was *Escherichia coli* K-12 MG1655 which has seven copies that fall into two clusters at 99% identity. SPAdes only assembled one version, whereas Matam assembled both (Emirge assembled none). However Matam also assembled more chimeras and fragments that clustered with the genuine *E. coli* sequences, and the sequences differed between the 100 bp and 150 bp libraries (Supplementary Figure 4). This suggests that reference-based assemblers like Emirge and Matam are more sensitive to the input read length or insert size, and also may produce spurious assemblies, especially with clusters of close sequences, and are hence less consistent.

When tested against the real-world environmental metagenomes, there was a strong linear relationship between the number of SSU rRNA reads extracted per library and the number of scaffolds (Emirge *R*^2^ = 0.683; Matam 0.558; SPAdes 0.669; *p* < 2×10^−16^ for all, Figure 5). Emirge assembled the most scaffolds per library (Emirge mean 16.9; Matam 6.95; SPAdes 7.48), but performs poorly by other metrics. Emirge scaffolds have more ambiguous bases (Emirge mean 2.775 Ns per scaffold; other assemblers 0), and are more likely to be chimeric (Emirge mean uchime_ref score 2.65; Matam 0.137; SPAdes 1.71). Matam scaffolds have the lowest chimerism scores, but are shorter (mean 1193 bp) than those from Emirge (1420 bp) or SPAdes (1368 bp). Matam scaffolds also have on average high identity to reference sequences in the database (Matam mean 99.8%; Emirge 97.7%; SPAdes 99.0%).

**Figure 5.**
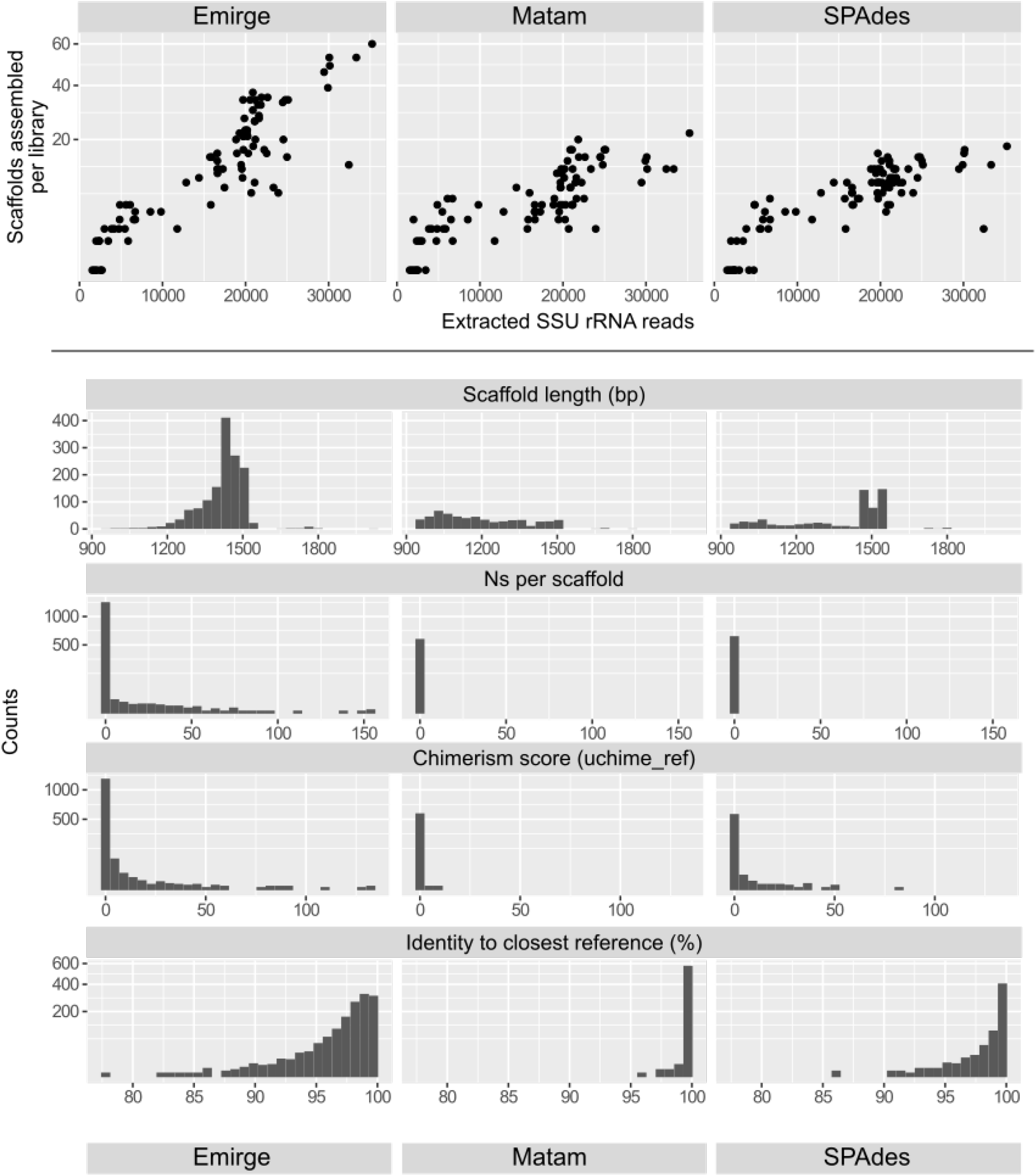
Comparison of scaffolds assembled by SPAdes, Emirge, and Matam using reads extracted by SortMeRNA (E-value 10^−5^, filtered reference database) from 85 Tara Oceans metagenomic libraries. (a) Plot of scaffolds assembled per library vs. number of extracted SSU rRNA reads. (b) Histograms of per-scaffold metrics for each assembly tool: scaffold length (bp), undetermined bases (Ns) per scaffold, uchime_ref chimerism score, and identity to closest reference sequence.

Chimeric sequences are an acknowledged problem and steps are taken to flag them in SILVA (28) but it is not possible to eliminate them entirely as the data ultimately comes from the wider scientific community. Chimera detection by reference-based methods, such as the uchime_ref score, are dependent on the reference database being free of chimeric sequences and having sufficiently close relatives to the query sequence. However, this cannot be guaranteed for public databases, where the deposited sequences have been assembled by different methods and tools, including reference-based assemblers that are known to produce chimeras, such as Emirge (16). Once chimeric sequences have been established in the reference database, subsequent chimerism scoring becomes less reliable. Based on our observed uchime_ref scores, some of the sequences assembled by either Emirge or SPADes may turn out to be chimeric, but each case will likely need to be examined separately.

Although Matam appears to be optimized for minimizing chimerism, this is at the expense of assembling more fragmentary scaffolds, and possibly overlooking SSU rRNA sequences that do not have a close relative in the reference database. SPAdes was therefore selected as the default in the phyloFlash pipeline as it yields a compromise between these two extremes, assembling longer scaffolds that are potentially more divergent as it is not dependent on a reference database.

### SSU rRNA-based metagenome analysis with phyloFlash

We designed phyloFlash (https://hrgv.github.io/phyloFlash/) for the rapid screening of SSU rRNA sequences in metagenomic libraries, through both a mapping-based overview of the taxonomic composition and full-length targeted assemblies. phyloFlash implements the database filtering described above, and allows the user to choose between different mapping tools (SortMeRNA or BBmap) and assemblers (Emirge and/or SPAdes), to evaluate the various trade-offs discussed above in a consistent way. The results and summary statistics are formatted as standalone HTML files, to make sharing and reporting easier. Additional utilities are also provided to compare the phyloFlash results from different libraries based on their taxonomic composition, e.g. graphically with bar plots or heat maps, or numerically as a distance matrix.

The default settings of phyloFlash have been chosen to allow quick run times on metagenomes of low to moderate diversity that have been sequenced as paired-end reads on Illumina platforms. High-diversity samples or those with uneven coverage, such as multiple displacement amplification libraries or metatranscriptomes, would likely need parameter optimizations.

Here, we present examples of how phyloFlash can be used to analyze a single metagenome, to compare multiple samples, and as part of a genome binning pipeline. Example commands to reproduce the phyloFlash runs are given in Supplementary Information.

#### Low-diversity metagenome and reference database completeness

We used a low-diversity, natural metagenome of known composition – a species of the marine catenulid flatworm *Paracatenula* sp. – to evaluate the effect of reference database completeness on the taxonomic composition and targeted assembly reported by the phyloFlash pipeline. Each species from this flatworm genus has an obligate symbiosis with a corresponding species of intracellular bacteria from the candidate genus Riegeria (Alphaproteobacteria: Rhodospirillaceae), which makes up ca. 40% of the biomass (41). The sequence library is hence predominantly composed of two genomes, one eukaryotic and one prokaryotic.

The majority of reads recovered by phyloFlash were either classified as Animalia (79.5%) or Rhodospirillales (17.8%), based on their best mapping hits to the SILVA reference database (Supplementary Figure 5). Each of the two assemblers included in the phyloFlash pipeline, SPAdes and Emirge, assembled full-length SSU rRNA sequences of both target organisms. This was expected because SSU rRNA sequences with >99% identity to the targets were already available in the SILVA database.

When SSU rRNA sequences >87% identical to those of the target organisms were removed from the reference database, which is equivalent to removing relatives up to the family level (21), targeted assembly could still yield a consistent picture of the metagenome composition, but reference-based classification of the extracted reads was less reliable. SPAdes assembled the same two full-length sequences as before, whereas Emirge only assembled a partial animal 18S rRNA sequence (962 bp, of which 203 were Ns), and no bacterial sequences (Supplementary Figure 6). The read classification based on mapping still assigned the majority of SSU rRNA reads to Animalia (75.5%) but only 1.5% to Rhodospirillales. Instead there was a higher proportion assigned to unclassified Alphaproteobacteria (9.6%), unclassified Proteobacteria (3.2%), or other alphaproteobacterial taxa (5.0%) (Supplementary Figure 5).

In comparison, the assembler Matam, which aligns contigs to the SSU rRNA reference database during scaffolding, produced fragmented assemblies when relatives were excluded from the reference database. Four sequences (508 to 740 bp) corresponded to the animal, and two (647 and 1043 bp) to the bacterium (Supplementary Figure 6). When the complete reference database was used, both the full-length target SSU rRNAs were assembled by Matam. This illustrates a possible drawback of reference-guided assembly approaches, such as those used by Emirge and Matam, in that their assemblies can be fragmented or incomplete when the target sequence is too divergent from existing reference sequences.

The lack of reference data from close relatives can also affect read classifiers that target whole-genome or protein-coding sequences. We could test the effect of reference data availability, as at the time of writing, Riegeria genome data were not yet publicly released. Furthermore, metagenomic read classifiers like Kaiju (13) and Kraken (14) include only microbial sequences as reference data but exclude animals and other multicellular eukaryotes. Kaiju assigned 17.4% of the reads to a taxonomic order, of which the most abundant taxa were Rhodospirillales (2.6%), fungal Glomerales (1.5%), and alphaproteobacterial Rhizobiales (0.88%) (Supplementary Figure 5). When only reads originating from the Riegeria genome were used as input, 14.7% could not be assigned to a taxonomic order, and 30.7% were classified as Rhodospirillales. At the class level, 55.0% were classified as Alphaproteobacteria.

SSU rRNA-based metagenome profiling therefore remains a powerful complement to read classifiers that use whole-genome data. Although the SSU rRNA makes up only a small percentage of total read data in most metagenomes, the reference databases for this gene are more phylogenetically representative than their equivalents for whole-genome data, particularly for eukaryotes. When samples contain divergent taxa whose close relatives are not represented in the reference database, taxonomic profiles based on SSU rRNA read classification may be misleading or inconsistent, and targeted assemblers that depend on alignment to a reference database may perform poorly, as shown in this example. Misclassification of reads could be exacerbated by inconsistencies between the taxonomy and the actual gene phylogeny (42). However, if they are sufficiently well-represented in the metagenome, full-length SSU rRNA sequences can still be assembled for divergent taxa by reference-independent assemblers, such as SPAdes.

#### Comparison of multiple metagenome samples

The taxonomic summary produced by the phyloFlash pipeline permits a rapid comparison of multiple metagenomes at a coarse-grained level, which can be followed by a finer-grained examination of specific taxa using the assembled sequences. To demonstrate the comparison of multiple samples, we applied the phyloFlash pipeline to metagenomes from six species of marine oligochaete worms of the genera *Olavius* and *Inanidrilus*. Each worm species in these genera has multiple species of symbiotic bacteria, with the most abundant symbiont being typically from the candidate genus Thiosymbion (Gammaproteobacteria: Arenicellales) (43). These worm species were chosen because their symbiont diversity is relatively low (no more than six bacteria per host species), and they have been characterized in previous studies by Sanger sequencing of SSU rRNA clone libraries and FISH (Supplementary Table 5) (44–49). Publicly-available sequencing data were directly downloaded and processed with phyloFlash using the script ENA_phyloFlash.pl that is provided with phyloFlash (Supplementary Information).

The results files from phyloFlash were directly used by the phyloFlash_compare.pl script to plot a synoptic overview of the read-mapping-based taxonomic composition for all samples, which can either be a heatmap (Figure 6) or barplot. Certain bacterial taxa occurred in multiple animal species and hence were likely to represent secondary symbionts, e.g. Desulfobacterales, Rhodospirillales, and Spirochaetales. The samples also fell into two clusters, one where Desulfobacterales and Spirochaetales were both present, and the other where Alphaproteobacteria (especially Rhodospirillales) were more abundant.

**Figure 6.**
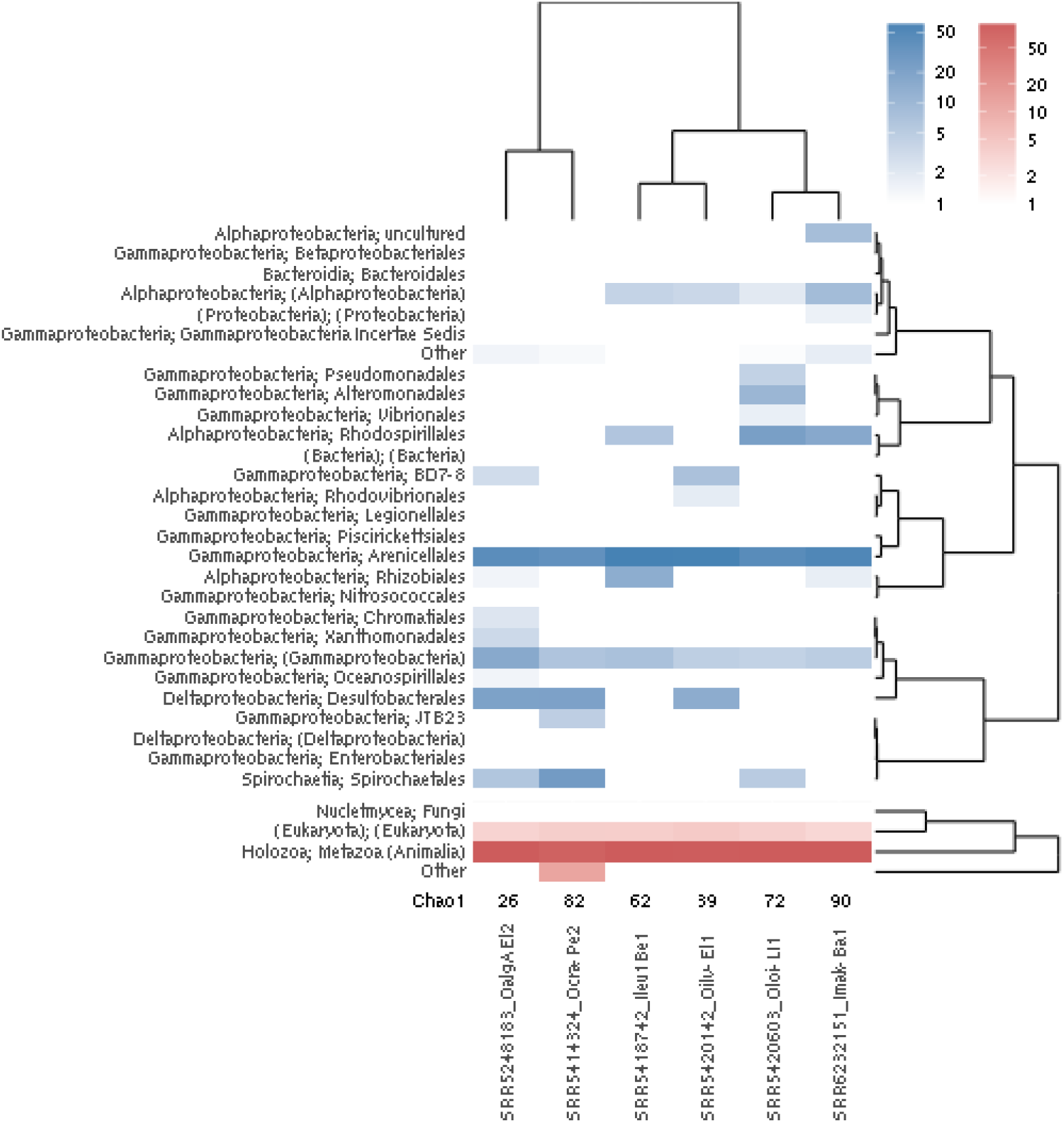
Heatmap of taxonomic assignments (rows) for SSU rRNA reads in metagenome libraries of six gutless oligochaete species with symbiotic consortia (columns). Plot was generated with the comparison scripts provided with phyloFlash. Color intensities represent the percentage of reads mapping to a given taxon, separated by prokaryotes (blue) and eukaryotes (red).

The full-length sequences produced by targeted assembly can give more specific information about the most abundant members of each metagenome, compared to the mapping-based taxonomic overview. Most of the previously-characterized symbiont SSU rRNA sequences were recovered from the metagenomes by targeted assembly with the phyloFlash pipeline, and those that were not recovered have been described as likely facultative symbionts (Supplementary Table 5). The metagenomes also contained bacterial SSU rRNA that were not previously reported from these host animals. Some had high sequence identity to known planktonic or sediment bacteria, e.g. *Vibrio tasmanensis* and *Psychrobacter* sp. from the *Olavius loisae* library, and hence may have been contamination from environmental bacteria. Each phyloFlash report file hyperlinks to GenBank records for the closest reference sequences to each assembled SSU rRNA sequence, for convenient browsing and comparison.

#### Assembly graph and SSU rRNA-based binning of microbial genomes

The SSU rRNA gene is an essential tool in microbial ecology, being used as a phylogenetic marker, and as a target for molecular probing to visualize specific taxa or species in environmental samples (19, 50). rRNA-targeted molecular ecology previously involved PCR amplification of the gene with specific primers, followed by one or more iterations of phylogenetic analysis, primer/probe design, microscopy, and PCR, the so-called rRNA cycle approach. Shotgun metagenomics can also yield full-length SSU rRNA sequences, assembled by phyloFlash or other tools, which can then be used as input into the “rRNA cycle”. In addition, microbial genomes can be “binned” from metagenome assemblies (metagenome-assembled genomes, or MAGs), offering functional information not otherwise accessible through the classical rRNA cycle.

However, MAGs often lack SSU rRNA sequences, as observed in several recent studies (1, 24). The SSU rRNA gene itself is difficult to assemble, because the gene has several highly conserved regions. These conserved regions lead to sets of shared k-mers among the SSU rRNA sequences present in the read pool that confound the assembly graph structure and eventually cause fragmented assemblies. Even if a full-length SSU rRNA gene is assembled, it may be difficult to assign to a specific MAG bin because the rRNAs often have multiple copies, that result in the rRNA genes being assembled into separate contigs from the rest of the genome. In addition, binning strategies often cannot assign such rRNA contigs to their correct bins because the rRNA genes are compositionally different. Coverage-based binning approaches also have similar problems, again due to the higher and variable copy number of rRNA genes. Although MAGs without SSU rRNA genes can still be phylogenetically informative and lead to the discovery of new microbial diversity (51), the lack of an SSU rRNA means that these genomes cannot be linked to known clades that have been defined on the basis of SSU rRNA sequences (21), nor can they be targeted by molecular probing for visualization or sorting (52).

To counter these problems, and to incorporate MAGs into the rRNA cycle of molecular ecology, we implemented a gene-targeted and assembly graph-based binning approach to assign rRNAs to genome bins. We used phyloFlash and two metagenome assemblers, Megahit (40) and MetaSPAdes (32), which both produce a graph-based representation of the assembly in the Fastg format (http://fastg.sourceforge.net/). This reports branching or circular connections between contigs, which can be exploited for genome binning in an automated and systematic fashion, but which would not be reported when the assembly is resolved into linear scaffolds. Each metagenome (Supplementary Table 6) was profiled with phyloFlash, and then assembled with Megahit and MetaSPAdes. Contigs connected to SSU rRNA sequences in the metagenome assembly graph were “fished” and compared to the SSU rRNA sequences from targeted assembly, with the script phyloFlash_fastgfishing.pl.

SSU rRNA-targeted binning worked reliably with low-diversity metagenomes that contained ten or fewer microbial genomes (Figure 7, Supplementary Table 6). As a first test case, we used the simulated metagenome which comprised ten genomes. Up to eight MAGs with high completeness and low contamination could be recovered.

**Figure 7.**
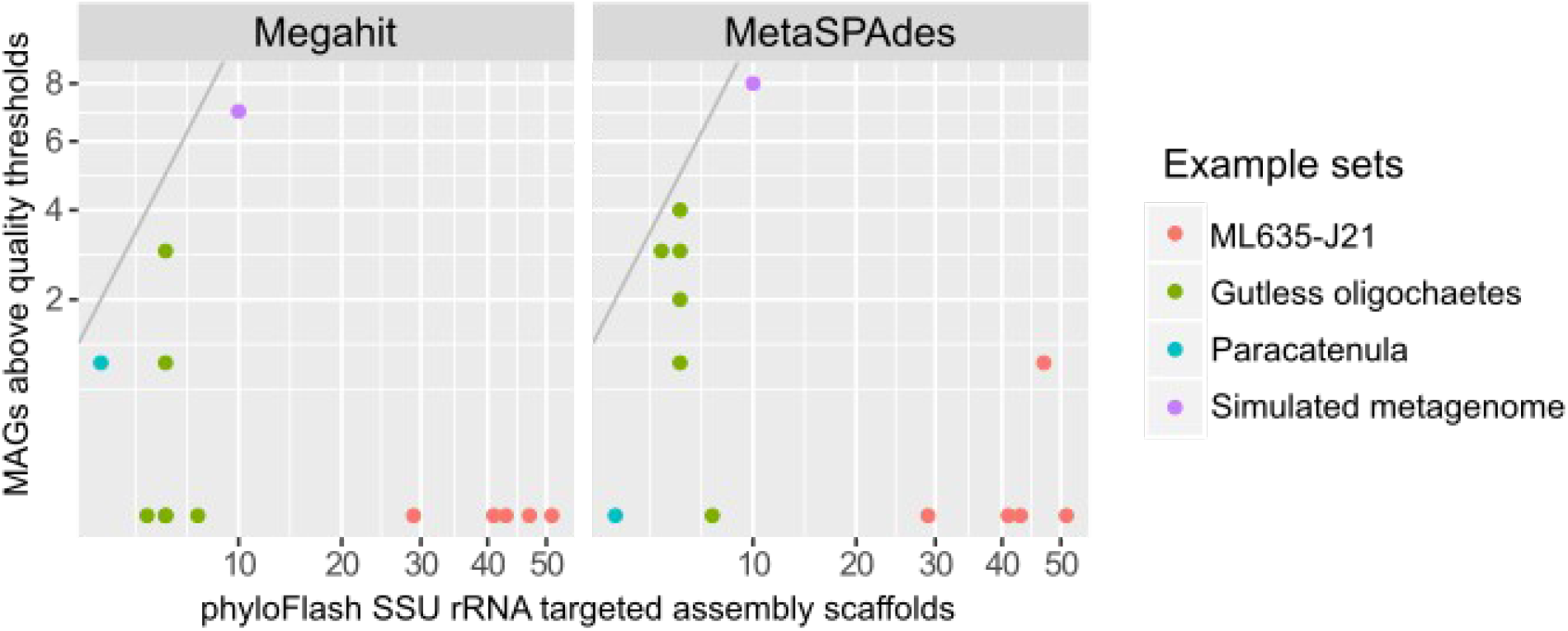
Genome recovery in assembly graph based binning depends on assembler software used. Comparison of MAGs above quality thresholds (> 70% completeness, < 10% contamination) that could be binned by SSU rRNA-targeted Fastg approach. The numbers of genome bins are plotted vs. SSU rRNA scaffold numbers assembled per library by SPAdes in phyloFlash pipeline. Reference line indicates 1:1 ratio.

We then moved to real-world examples, starting again with the simple monospecific *Paracatenula* metagenome. Both assemblers produced graphs that yielded bins, but at different completeness. Even though the input read set only had a mean nucleotide coverage of 39.3, the recovered bin from the Megahit assembly was a complete representation of the symbiont’s genome. Manual inspection of the assembly graph using Bandage showed a closed bi-circular representation, with the rRNA operon forming the only repeat region. The CheckM pipeline underestimated genome completeness, as it belongs to an endosymbiont that has experienced genome reduction (Jaeckle et al., submitted). In contrast to the complete microbial genome recovered from the Megahit assembly, only half the genome was recovered from the MetaSPAdes assembly graph. This could be due to the low coverage of the target genome of 1–3x k-mer coverage at the longest k-mer employed and to different levels of low-count k-mer removal in the assembly pipelines.

In a next step, we tested the targeted binning approach on the microbial communities of the gutless oligochaetes. More than half of the microbial community from the gutless oligochaetes predicted by phyloFlash targeted SSU rRNA assemblies (between four and seven per library, Supplementary Table 6) could be recovered as MAGs of moderate to high quality (> 70% completeness, < 10% contamination). We again observed different levels of completeness from the two assemblers. Unlike the *Paracatenula* dataset, MetaSPAdes yielded consistently better results than Megahit, likely due to the higher coverage of each genome compared to the *Paracatenula* dataset.

We expected that the performance of our assembly graph-based genome binning procedure would deteriorate as the microbial community, and hence the assembly graph, becomes more complex. This would happen when different genomes contain SSU rRNA sequences that share enough k-mers to prevent the unambiguous assembly of full-length genes, resulting either in fragmented or linked assemblies. Furthermore, many genomes in a high-diversity metagenome may not have adequate coverage to permit a linked assembly graph and therefore the binning of a complete MAG, as observed in the low-coverage *Paracatenula* example. However, these drawbacks can be turned into an advantage when specific organisms are being targeted for assembly. By screening metagenomic read sets using phyloFlash for full-length assemblies of the SSU rRNA of a given target species or taxon, one could select which read sets to assemble. Such prior screening could drastically reduce the computational load for recovering high quality genome bins for a defined group of organisms, and can be easily integrated into existing assembly and binning pipelines.

To test the screening of metagenomes by phyloFlash for specific organisms of interest, we focused on an enigmatic bacterial clade called Marinamargulisbacteria (Carnevali et al., BioRxiv: https://doi.org/10.1101/328856). The first genomes of Marinamargulisbacteria were recently reported under the name UBP8, among sixteen other novel phylum-level clades (UBP1 to 17) that were recovered in a set of over 8000 MAGs binned from a wide collection of metagenomic samples (1). Marinamargulisbacteria belongs to the Margulisbacteria, a clade that is sister to Cyanobacteria. Marinamargulisbacteria was initially designated as a separate phylum, but is currently classified as a class of Margulisbacteria and is listed as ‘GWF2-35-9’ in the GTDB (53). Carnevali et al. designated the name Marinamargulisbacteria because all members come from marine samples, and they also provided new observations from single cell amplified genomes (SAGs). Currently, one MAG and five SAGs of poor to moderate quality are available for UBP8/Marinimargulisbacteria. None of the initially recovered MAGs had a full-length SSU rRNA gene; only one featured a 500 bp fragment that was tentatively annotated as belonging to a clade designated ML635J-21 (1). To link full-length SSU rRNA information to the Marinamargulisbacteria, we set up a targeted reanalysis of all Tara Oceans samples reported to contain Marinamargulisbacteria/UBP8. Using phyloFlash, we recovered full length SSU rRNA that fell into ML635J-21/Margulisbacteria (SILVA 128/132 taxonomy) from all of these samples. We then applied our SSU rRNA-targeted binning approach via assembly graph-based fishing, and recovered a complete MAG based on the closed graph structure from sample ERR59434 (98% completeness and 1.72% contamination, of which 100% can be explained by strain variation). Based on phylogenomic analyses using 43 conserved marker genes, the genome is the complete MAG representation of a population genome that had previously been recovered at 51.72% completeness with traditional binning approaches (accession DBHN01.1) (1).

In practice, which assembler and binning software produces the best results for a given dataset has to be determined empirically. phyloFlash aids this process by giving a quick overview of the expected diversity in the library, and with its command line based binning procedure that exploits the assembly graph as well as the properties of the SSU gene. phyloFlash can be effectively used to evaluate and compare different assemblers and assembly parameters and can complement existing binning strategies (23).

### Conclusion

The SSU rRNA gene remains indispensable to molecular ecology in the current era of whole-genome metagenomics, particularly because of the extensive reference data that has been accumulated for this gene, and the use of rRNA as a target for molecular probing. phyloFlash is a complete workflow for SSU rRNA-targeted profiling of metagenomes that integrates database preparation, read extraction, taxonomic classification, targeted assembly and SSU rRNA targeted binning. It has been designed to provide a quick and user-friendly overview of the composition of metagenomic sequence libraries based on all domains of life, and can be used for the rapid screening and comparison of multiple samples, and for retrieving complete SSU rRNA genes and the associated genomic bins.

## AVAILABILITY

phyloFlash is available at https://github.com/HRGV/phyloFlash under a GNU GPL3 license. The software is also archived on Zenodo at doi:10.5281/zenodo.1327399. The version described in this paper is v3.3. Results from the usage examples have been archived at doi:10.5281/zenodo.1464895.

## ACKNOWLEDGEMENT

We thank our colleagues and phyloFlash users who have contributed to the software development by reporting bugs and suggesting new features. We also thank Nicole Dubilier for arranging financial support, and Antonio Fernandez-Guerra for help with accessing test data. Oligochaete metagenome data in the usage examples were produced at the US Department of Energy Joint Genome Institute in collaboration with the Max Planck Institute for Marine Microbiology under the Community Science Program.

## AUTHOR CONTRIBUTIONS

H.G.V. conceived project and wrote initial version of software. E.P. and B.S. developed software. B.S. and H.G.V. tested software and analyzed data. B.S. and H.G.V. wrote manuscript.

