## Supplementary Information for "phyloFlash – Rapid SSU rRNA profiling and targeted assembly from metagenomes"

**Supplementary Information.** Code to reproduce phyloFlash usage examples

The code presented (for Unix/Linux systems) assumes that phyloFlash scripts are accessible from the path, dependencies have already been installed, and that the user is not behind a proxy server. Each analysis should be run in a separate folder. Detailed installation and usage instructions are provided in the online manual: https://hrgv.github.io/phyloFlash/

### Download and set up phyloFlash database

```
phyloFlash_makedb.pl --remote
```

### Low-diversity metagenome

```
ENA_phyloFlash.pl --acc ERR2931548 -phyloFlash="--everything --readlength 150" --CPUs 12
```

### Comparison of multiple metagenome samples

```
## Download reads from ENA by run accession number
## and run phyloFlash with a single wrapper script
for ACC in SRR5248183 SRR5413717 SRR5418742 SRR5420142 SRR5420603 SRR6232151
do
ENA_phyloFlash.pl --acc $ACC --phyloFlash="--everything" --CPUs 12
done

## Compare samples and plot heatmap in PDF format
phyloFlash_compare.pl --task heatmap --allzip --level 4 \
--out phyloFlash_gutless_oligochaete_comparison --outfmt pdf
```

### SSU rRNA-based binning of microbial genomes

```
LIB=ERR594304 # Library name prefix
READF=ERR594304_1.fastq.gz # Forward read file
READR=ERR594304_2.fastq.gz # Reverse read file

#run phyloFlash on raw reads
#phyloFlash.pl -lib ${LIB}_pf -read1 $READF -read2 $READF \
-everything -CPUs 12 -readlength 100

#quality and adapter trim reads
mkdir  trimmed
cd trimmed
bbduk.sh ref=adapters.fa ktrim=l minlength=36 mink=11 hdist=1 \
in=../$READF in2=../$READR out=${LIB}_ktriml.fq.gz
bbduk.sh ref=adapters.fa ktrim=r trimq=2 qtrim=rl minlength=36 mink=11 hdist=1 \
in=${LIB}_ktriml.fq.gz interleaved=t out=${LIB}_q2_ktrimmed.fq.gz
cd ..
#kmer filter reads
mkdir filtered/
cd filtered
bbnorm.sh -Xmx200g in=../trimmed/${LIB}_q2_ktrimmed.fq.gz \
lowbindepth=3 highbindepth=6 outhigh=${LIB}_q2_ktrimmed_k31higher6.fq.gz \
passes=1 threads=24 interleaved=t
cd ..

#assemble with megahit
as=${LIB}_all_kfilt_MH_21_93; #rename
mkdir assemblies -p
cd assemblies
megahit --k-min 21 --k-max 93 --k-step 20 -m 0.9 -t 48 \
--out-prefix $as \
--12 ../filtered/${LIB}_q2_ktrimmed_k31higher6.fq.gz \
-o $as
megahit_toolkit contig2fastg 93 ./${as}/intermediate_contigs/k93.contigs.fa > ./${as}/${as}.fastg
cd ..

#fastgfish megahit assembly
mkdir bins -p
cd bins
ln -s ../assemblies/${as}/intermediate_contigs/k93.contigs.fa \
./${as}.k93.contigs.fasta
```

```
ln -s ../assemblies/${as}/${as}.fastg .
phyloFlash_fastgFishing.pl --fasta ${as}.k93.contigs.fasta \
--fastg ${as}.fastg \
--out ${as}_fastgbin --compare-zip \
../${LIB}_pf.phyloFlash.tar.gz \
--assembler megahit --outfasta \
--min-SSU-frac 0.6

cd ../assemblies

#assemble with spades
as_sp=${LIB}_all_kfilt_SPm_21_93; #rename
spades.py --meta -k 21,33,55,77,93 -m 880 -t 48 \
-o $as_sp --12 ../filtered/${LIB}_q2_ktrimmed_k31higher6.fq.gz

#fastgfish spades assembly
cd ../bins;
ln -s ../assemblies/${as_sp}/scaffolds.fasta ./${as_sp}.scaffolds.fasta
ln -s ../assemblies/${as_sp}/assembly_graph.fastg \
./${as_sp}.assembly_graph.fastg
ln -s ../assemblies/${as_sp}/scaffolds.paths \
./${as_sp}.scaffolds.paths
phyloFlash_fastgFishing.pl --fasta ${as_sp}.scaffolds.fasta \
--fastg ${as_sp}.assembly_graph.fastg \
--paths ${as_sp}.scaffolds.paths \
--out ${as_sp}_fastgbin \
--compare-zip ../${LIB}_pf.phyloFlash.tar.gz \
--assembler spades --outfasta --min-SSU-frac 0.6
```

### Supplementary Files

The following data have been deposited in Zenodo under doi:10.5281/zenodo.1464895

| | |
|---|---|
| tara_oceans_example/ | phyloFlash and Matam runs for 85 Tara Oceans metagenomes (10 M read subsets) |
| phyloFlash_usage_example_gutless_oligochaetes/ | phyloFlash comparison script usage example for six gutless oligochaete metagenome libraries |
| phyloFlash_usage_example_Paracatenula/ | phyloFlash, Matam, and Kaiju results for Paracatenula flatworm metagenome using reference databases with and without the target organisms |
| simulated_metagenome_100bp/ | phyloFlash and Matam results from simulated metagenome (100 bp reads), and code to generate the reads. |
| simulated_metagenome_150bp/ | phyloFlash and Matam results from simulated metagenome (150 bp reads), and code to generate the reads. |
| fastg_fishing_genome_bins/ | Metagenome assemblies for usage examples with MAGs from SSU rRNA-targeted binning approach |

**Supplementary Table 1**. RefSeq genomes used for simulated metagenome library.

| Accesssion | Description | No. SSU rRNA genes |
|---|---|---|
| NC_000918.1 | *Aquifex aeolicus* VF5 chromosome, complete genome | 2 |
| NC_004307.2 | *Bifidobacterium longum* NCC2705 chromosome, complete genome | 4 |
| NC_002528.1 | *Buchnera aphidicola* str. APS (Acyrthosiphon pisum) chromosome, complete genome | 1 |
| NC_002971.4 | *Coxiella burnetii* RSA 493, complete genome | 1 |
| NC_000913.3 | *Escherichia coli* str. K-12 substr. MG1655, complete genome | 7 |
| NC_007644.1 | *Moorella thermoacetica* ATCC 39073 chromosome, complete genome | 1 |
| NC_005042.1 | *Prochlorococcus marinus* subsp. *marinus* str. CCMP1375 complete genome | 1 |
| NC_007643.1 | *Rhodospirillum rubrum* ATCC 11170 chromosome, complete genome | 4 |
| NC_007677.1 | *Salinibacter ruber* DSM 13855 chromosome, complete genome | 1 |
| NC_009636.1 | *Sinorhizobium medicae* WSM419 chromosome, complete genome | 3 |

**Supplementary Table 2**. Read extraction statistics for simulated metagenomes, comparing different sequence library profiles, extraction software, and stringency settings.

| Library profile | Software | Setting | Reads extracted | | Total | Selectivity | Sensitivity | Timing (min) |
|---|---|---|---|---|---|---|---|---|
| | | | Non-SSU rRNA | SSU rRNA | | | | |
| HiSeq2000, 100 bp PE | BBmap | Identity > 50% | 4349 | 19473 | 23822 | 0.817 | 0.963 | 2.35 |
| HiSeq2000, 100 bp PE | BBmap | Identity > 60% | 3516 | 19420 | 22936 | 0.847 | 0.961 | 2.40 |
| HiSeq2000, 100 bp PE | BBmap | Identity > 70% | 1747 | 19021 | 20768 | 0.916 | 0.941 | 2.75 |
| HiSeq2000, 100 bp PE | SortMeRNA | E-value < $10^{-5}$ | 4126 | 19245 | 23371 | 0.823 | 0.952 | 11.78 |
| HiSeq2000, 100 bp PE | SortMeRNA | E-value < $10^{-7}$ | 3222 | 19143 | 22365 | 0.856 | 0.947 | 9.40 |
| HiSeq2000, 100 bp PE | SortMeRNA | E-value < $10^{-9}$ | 2392 | 19035 | 21427 | 0.888 | 0.942 | 9.02 |
| HiSeq2500, 150 bp PE | BBmap | Identity > 50% | 2093 | 13246 | 15339 | 0.864 | 0.958 | 1.38 |
| HiSeq2500, 150 bp PE | BBmap | Identity > 60% | 1489 | 13183 | 14672 | 0.899 | 0.953 | 1.38 |
| HiSeq2500, 150 bp PE | BBmap | Identity > 70% | 650 | 12868 | 13518 | 0.952 | 0.931 | 1.43 |
| HiSeq2500, 150 bp PE | SortMeRNA | E-value < $10^{-5}$ | 4902 | 13368 | 18270 | 0.732 | 0.967 | 13.7 |
| HiSeq2500, 150 bp PE | SortMeRNA | E-value < $10^{-7}$ | 3809 | 13331 | 17140 | 0.778 | 0.964 | 13.4 |
| HiSeq2500, 150 bp PE | SortMeRNA | E-value < $10^{-9}$ | 2990 | 13288 | 16278 | 0.816 | 0.961 | 13.4 |

**Supplementary Table 3**. Accession numbers of marine planktonic metagenomes from the Tara Oceans project used for comparison of different mapping and assembly tools, testing of database filtering on read extraction, and SSU-rRNA-targeted genome binning usage example.

| Accession number | Read extraction and targeted assembly comparison | Database filtering and read extraction example | SSU rRNA-targeted genome binning example |
|---|---|---|---|
| ERR315856 | + | + | |
| ERR315857 | + | + | |
| ERR315858 | + | + | |
| ERR315859 | + | + | |
| ERR315860 | + | + | |
| ERR315861 | + | | |
| ERR315862 | + | | |
| ERR315863 | + | | |
| ERR318618 | + | | |
| ERR318620 | + | | |
| ERR588857 | + | | |
| ERR594284 | + | | |
| ERR594285 | + | | |
| ERR594286 | + | | |
| ERR594287 | + | | |
| ERR594288 | + | | |
| ERR594289 | + | | |
| ERR594290 | + | | |
| ERR594291 | + | | |
| ERR594292 | + | | |
| ERR594293 | + | | |
| ERR594294 | + | | |
| ERR594295 | + | | |
| ERR594296 | + | | |
| ERR594297 | + | | |
| ERR594298 | + | | |
| ERR594299 | + | | |
| ERR594300 | + | | |
| ERR594301 | + | | |
| ERR594302 | + | | |
| ERR594303 | + | | |
| ERR594304 | + | | + |
| ERR594305 | + | | |
| ERR594306 | + | | |
| ERR594307 | + | | |
| ERR594308 | + | | |
| ERR594309 | + | | |
| ERR594310 | + | | |
| ERR594311 | + | | + |
| ERR594312 | + | | |
| ERR594313 | + | | + |
| ERR594314 | + | | |

| | | | |
|---|---|---|---|
| ERR594315 | + | | |
| ERR594316 | + | | |
| ERR594317 | + | | |
| ERR594318 | + | | |
| ERR594319 | + | | |
| ERR594320 | + | | |
| ERR594321 | + | | |
| ERR594322 | + | | |
| ERR594323 | + | | |
| ERR594324 | + | | |
| ERR594325 | + | | + |
| ERR594326 | + | | |
| ERR594327 | + | | |
| ERR594328 | + | | |
| ERR594329 | + | | |
| ERR594330 | + | | |
| ERR594331 | + | | |
| ERR594332 | + | | |
| ERR594333 | + | | |
| ERR594334 | + | | |
| ERR594335 | + | | |
| ERR594336 | + | | |
| ERR594337 | + | | |
| ERR594338 | + | | |
| ERR594339 | + | | |
| ERR594340 | + | | |
| ERR594341 | + | | |
| ERR594342 | + | | + |
| ERR594343 | + | | |
| ERR594344 | + | | |
| ERR594345 | + | | |
| ERR594346 | + | | |
| ERR594347 | + | | |
| ERR594348 | + | | |
| ERR594349 | + | | |
| ERR594352 | + | | |
| ERR594353 | + | + | |
| ERR594354 | + | + | |
| ERR594355 | + | + | |
| ERR594356 | + | + | |
| ERR594357 | + | + | |
| ERR594358 | + | + | |
| ERR594359 | + | + | |

**Supplementary Table 4**. Ratio of assembled reads extracted by BBmap vs. SortMeRNA for different software settings, from 85 Tara Oceans metagenomic libraries (four libraries yielded no assembled reads).

| SortMeRNA E-value | BBmap min. %id | Min. | 1st Quartile | Median | Mean | 3rd Quartile | Max. |
|---|---|---|---|---|---|---|---|
| $10^{-5}$ | 50 | 0 | 0.840 | 0.934 | 0.882 | 0.988 | 1.11 |
| $10^{-5}$ | 60 | 0 | 0.816 | 0.923 | 0.869 | 0.991 | 1.11 |
| $10^{-5}$ | 70 | 0 | 0.802 | 0.918 | 0.859 | 0.992 | 1.13 |
| $10^{-7}$ | 50 | 0 | 0.840 | 0.933 | 0.895 | 0.998 | 1.26 |
| $10^{-7}$ | 60 | 0 | 0.822 | 0.922 | 0.881 | 0.997 | 1.12 |
| $10^{-7}$ | 70 | 0 | 0.810 | 0.926 | 0.870 | 0.999 | 1.12 |
| $10^{-9}$ | 50 | 0 | 0.853 | 0.934 | 0.906 | 1.000 | 1.63 |
| $10^{-9}$ | 60 | 0 | 0.843 | 0.928 | 0.888 | 0.998 | 1.04 |
| $10^{-9}$ | 70 | 0 | 0.816 | 0.928 | 0.878 | 1.00 | 1.04 |

**Supplementary Table 5**. Gutless oligochaete species used for microbiome comparison example.

| Accession | Species | Symbionts per individual previously described | Symbiont SSU rRNA assembled from metagenome | Other bacterial SSU rRNA assembled from metagenome | Description of symbionts |
|---|---|---|---|---|---|
| SRR5248183 | Olavius algarvensis | 3 to 5 | 4 | 0 | (Dubilier et al., 2001; Kleiner et al., 2012) |
| SRR5414324 | Olavius crassitunicatus | 3 to 6 | 5 | 0 | (Blazejak et al., 2005) |
| SRR5418742 | Inanidrilus leukodermatus | 3 to 5 | 3 | 0 | (Blazejak et al., 2006) |
| SRR5420142 | Olavius ilvae | 3 to 4 | 3 | 1 | (Ruehland et al., 2008) |
| SRR5420603 | Olavius loisae | 3 | 3 | 3 | (Dubilier et al., 1999) |
| SRR6232151 | Inanidrilus makropetalos | 3 | 3 | 1 | (Blazejak et al., 2006) |

**Supplementary Table 6**. Comparison of MAGs above quality thresholds (> 70% completeness, < 10% contamination) that could be binned by SSU rRNA-targeted Fastg approach vs. SSU rRNA scaffold numbers assembled in phyloFlash pipeline per library (including Eukaryota).

| **Example set** | **Accession** | **phyloFlash targeted SSU rRNA assemblies** | | **MAGs above quality thresholds** | |
|---|---|---|---|---|---|
| | | **Emirge** | **SPAdes** | **Megahit** | **MetaSPAdes** |
| ML635J-21 | ERR594304 | 179 | 47 | 0 | 1 |
| ML635J-21 | ERR594311 | 202 | 43 | 0 | 0 |
| ML635J-21 | ERR594313 | 38 | 29 | 0 | 0 |
| ML635J-21 | ERR594325 | 183 | 51 | 0 | 0 |
| ML635J-21 | ERR594342 | 2 | 41 | 0 | 0 |
| Gutless oligochaete | SRR5248183 | 7 | 5 | 1 | 1 |
| Gutless oligochaete | SRR5414324 | 7 | 5 | 0 | 2 |
| Gutless oligochaete | SRR5418742 | 7 | 4 | 0 | 3 |
| Gutless oligochaete | SRR5420142 | 5 | 5 | 0 | 4 |
| Gutless oligochaete | SRR5420603 | 5 | 7 | 0 | 0 |
| Gutless oligochaete | SRR6232151 | 5 | 5 | 3 | 3 |
| *Paracatenula* | ERR2931548 | 2 | 2 | 1 | 0 |
| Simulated metagenome | (NA) | 9 | 10 | 7 | 8 |

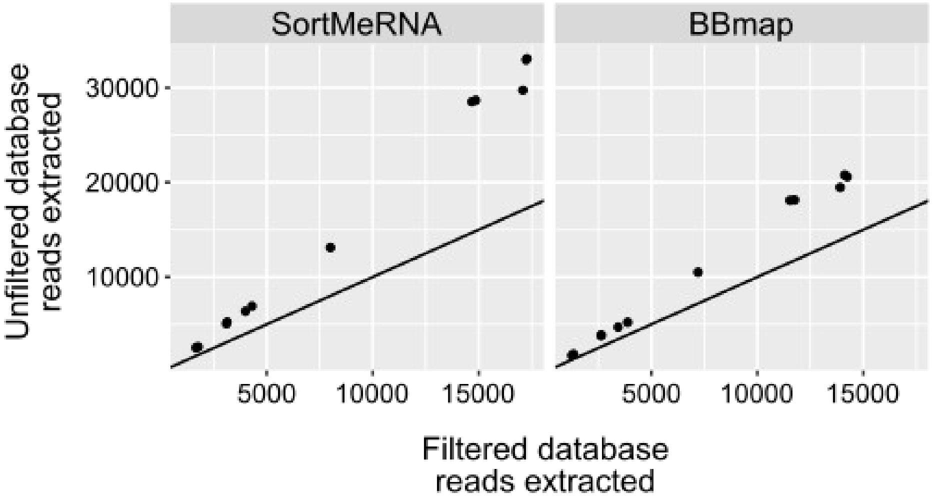


**Supplementary Figure 1**. Plots of total reads extracted by SortMeRNA (left) or BBmap (right) using unfiltered vs. filtered reference database; the 1:1 line is overlain for comparison.

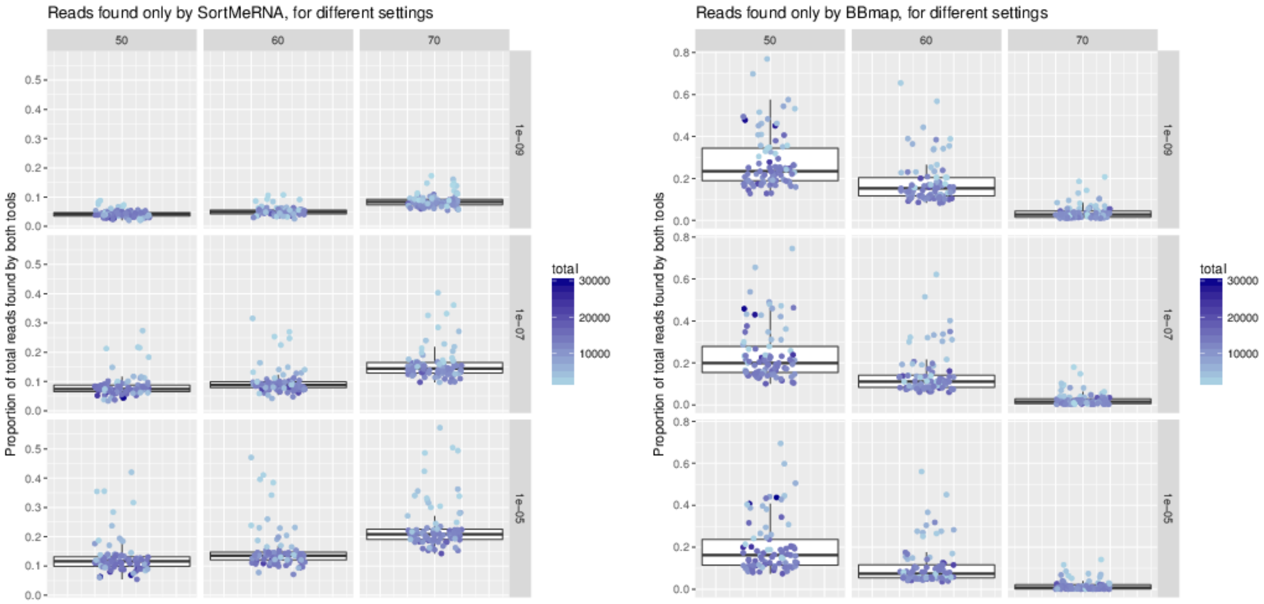


**Supplementary Figure 2**. Boxplots of the fraction of reads extracted by only SortMeRNA (left) or only BBmap (right) from Tara Oceans metagenome libraries, comparing different cutoff settings for BBmap (horizontal facets) and SortMeRNA (vertical facets).

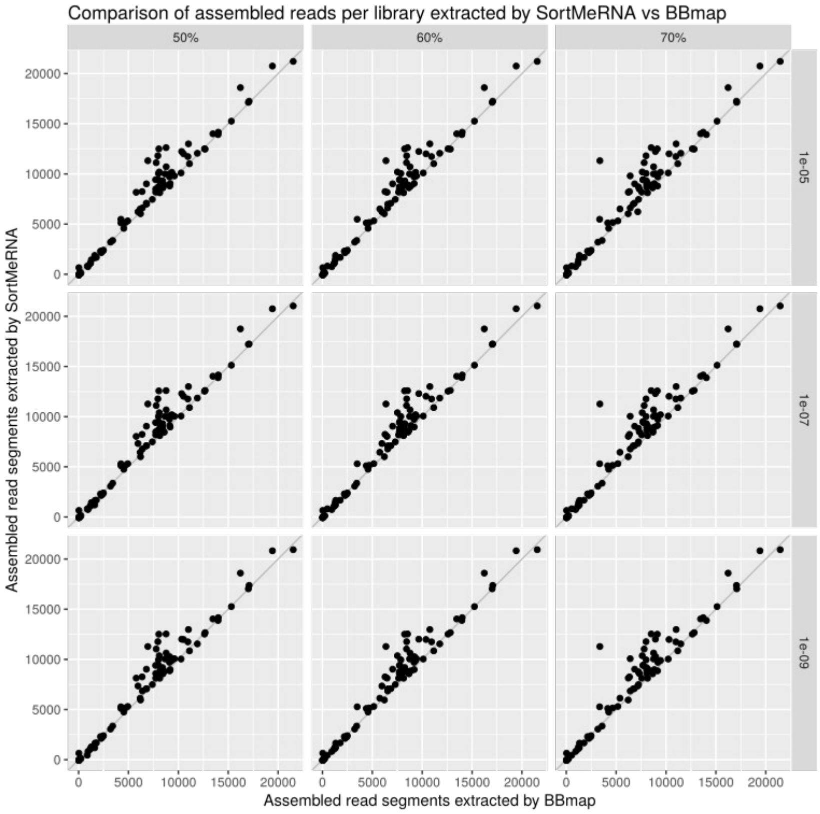


**Supplementary Figure 3**. Numbers of reads assembled to full-length sequences per library that were extracted by SortMeRNA (vertical axes) vs. BBmap (horizontal axes), comparing different cutoff settings for SortMeRNA (vertical facets) vs. BBmap (horizontal facets). The 1:1 line (grey) is added to each plot for reference.

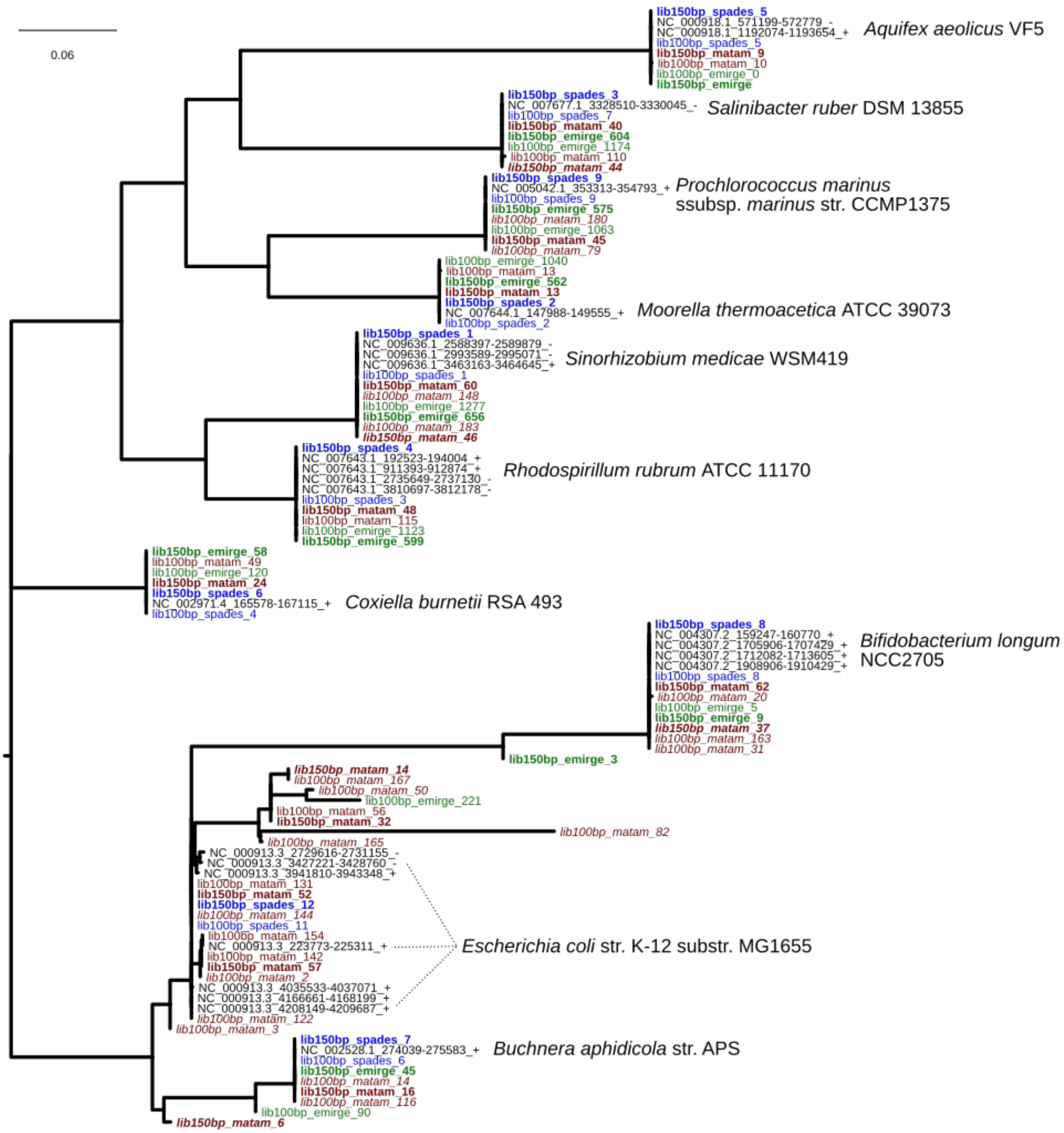


**Supplementary Figure 4**. Phylogenetic tree of SSU rRNA sequences assembled by Matam (red), Emirge (green), and SPAdes (blue) from simulated metagenome libraries, 100 bp paired-end (normal weight) and 150 bp paired-end (bold), alongside the original gene sequences. *Italics* – Sequences shorter than 60% of full-length SSU rRNA.

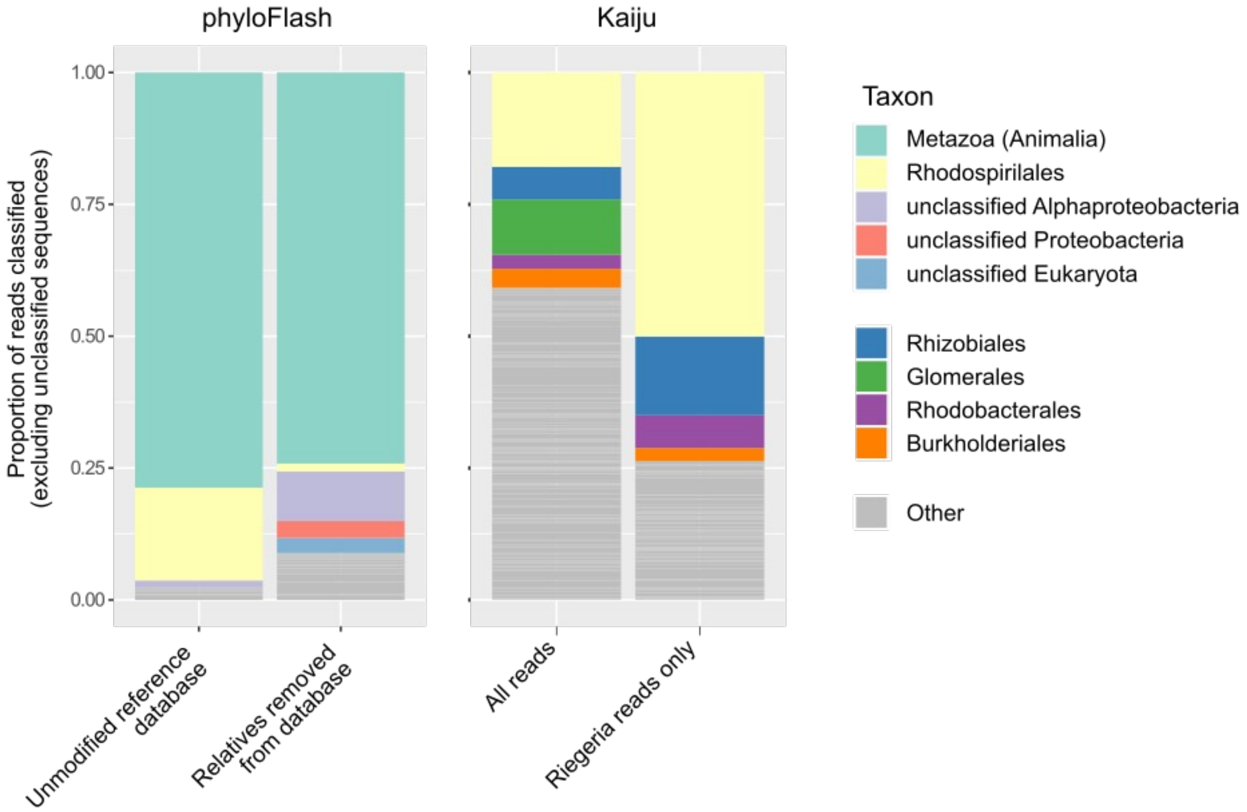


**Supplementary Figure 5**. Taxonomic composition (at order level) of a low-diversity platyhelminth metagenome. Composition based on SSU rRNA reads via phyloFlash pipeline (left), before and after sequences related to the known target organisms were removed from the reference database. Compared to the composition predicted with protein-coding sequences via the Kaiju pipeline (right), excluding reads with no assignment or which could not be classified to a taxonomic order.

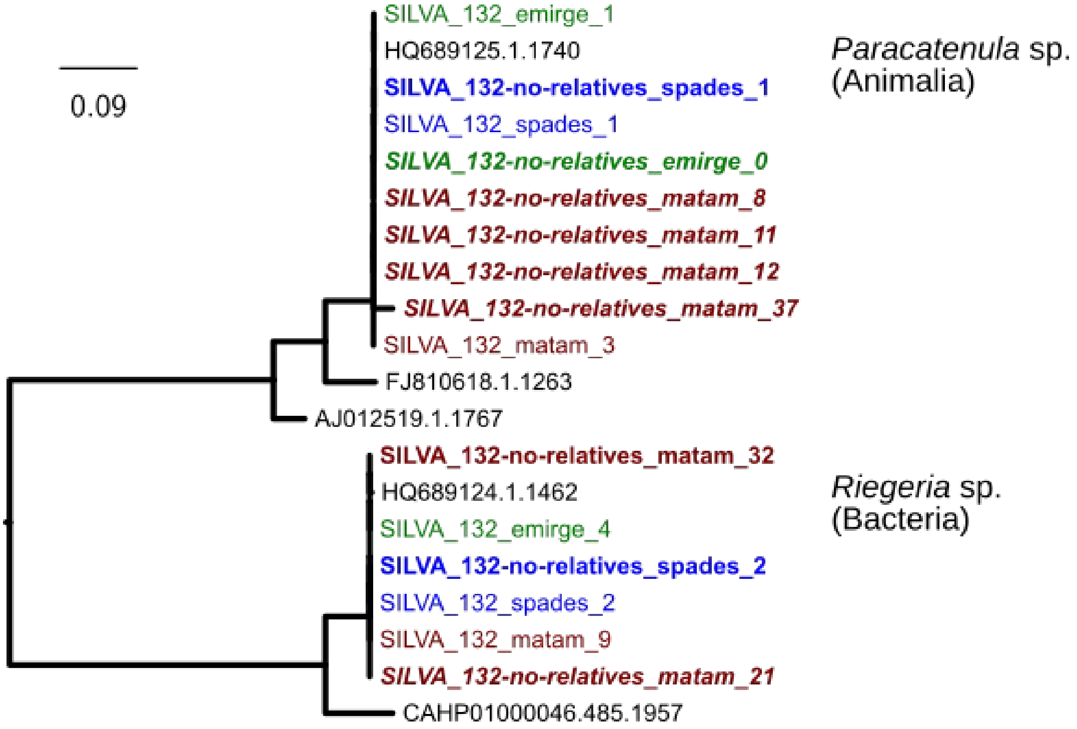


**Supplementary Figure 6**. Phylogenetic tree of SSU rRNA sequences assembled by Matam (red), Emirge (green), and SPAdes (blue) from a low-diversity platyhelminth metagenome, using either the SILVA 132 SSU Ref NR96 database as reference for read extraction and assembly (normal weight), or with sequences >87% identical to the target organisms removed from the reference database (bold weight). *Italics* - sequences <60% of full-length SSU rRNA.
